# Exploring the Dynamics of Nonlinear Biochemical Systems using Control-Based Continuation

**DOI:** 10.1101/695866

**Authors:** Brandon Gomes, Irene de Cesare, Agostino Guarino, Mario di Bernardo, Ludovic Renson, Lucia Marucci

## Abstract

Mathematical modelling is routinely used in Systems Biology to understand the mechanisms causing nonlinear phenomena in gene expression, such as switch-like behaviours and temporal oscillations. The reliability of model predictions and bifurcation analysis depend on modelling assumptions and specific choices of model parameters; however, the identification of models is highly challenging due to the complexity of biochemical interactions and noise in experimental data.

This paper numerically investigates the use of control-based continuation (CBC) for tracking dynamical features of biochemical systems and, in particular, the bistable dynamics of a gene regulating pluripotency in embryonic stem cells.

CBC is a method that exploits feedback control and path following algorithms to explore the dynamic features of a nonlinear physical system directly during experimental tests. CBC applications have so far been limited to non-living (i.e. electro-mechanical) systems. Our numerical simulations show that, in principle, CBC could also be applied to biological experiments to characterise the switch-like dynamics of genes that are important for cell decision making.

## 1 INTRODUCTION

Complex nonlinear dynamics naturally emerges in biological systems; in particular, feedback loops in gene regulatory networks (GRNs), both in natural and engineered systems, can cause the emergence of switch-like or oscillatory gene expression dynamics, which, in turn, can regulate important processes in cells and organisms, such as differentiation and development [1–5]. A combination of experiments and mathematical modelling is generally used in order to study nonlinear dynamics in gene expression and other dynamical properties of GRNs.

Models are used to generate predictions about the system dynamics and to perform bifurcation analysis via numerical continuation, which allows the tracking of solutions while certain parameters change using path-following techniques (e.g. pseudo-arclength continuation [6, 7]).

The identification of biochemical models is highly challenging across applications [8] and model structure can be non-unique for a given biochemical system and corresponding experimental data [9]. As such, parameter and model uncertainty can significantly alter the reliability of model predictions and bifurcation studies. As of now, there exists no method that can directly measure nonlinear dynamic features such as bifurcations by using biological experiments.

Control-Based Continuation (CBC) is a non-parametric method that maps out the dynamic features of a nonlinear physical system directly during experimental tests, without relying on a mathematical model [10]. Combining feedback control with numerical continuation algorithms, CBC modifies on-line the input applied to the system in order to isolate the nonlinear behaviours of interest. In this way, CBC offers the best conditions to analyse dynamic features in detail, to follow them as experimentally-controllable parameters are changed, and to detect and track boundaries between qualitatively different types of behaviour (i.e. bifurcations). The fundamental principles of CBC are well established, and the method has been applied to a wide range of non-living (i.e. electro-mechanical) systems: for instance, a parametrically excited pendulum [10] and a bilinear oscillator [11]. We recently used CBC on a multi-degree-of-freedom structure exhibiting complex nonlinear phenomena such as mode interaction, quasi-periodic oscillations and isolated response curves [12,13].

Here, building on existing CBC methodologies [21], we first track the solutions of the normal form of a fold bifurcation. Then, we use CBC to track stationary solutions of Nanog, a gene regulating mouse embryonic stem cell pluripotency. Nanog bistability has been extensively reported experimentally [15–18], and various mathematical models have been proposed to explain such heterogeneity [9]. A mathematical formalism we recently developed [19] is exploited here to simulate the network dynamics. Our results demonstrate the versatility of the CBC method whose experimental implementation in living cells could open new research avenues to understand nonlinear biochemical system dynamics.

## 2 RESULTS

### 2.1 Principles of Control-Based Continuation

The fundamental principles of the CBC method used in this paper are discussed using the scalar differential equation

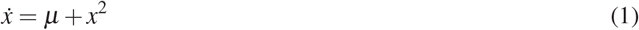

where *x* ∈ ℝ is the system’s state and *μ* ∈ ℝ is the bifurcation parameter. Eq. (1) corresponds to the normal form of a fold (or saddle-node) bifurcation. The equilibria of the above system are given by the formula:

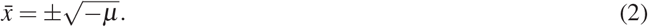

Their stability can be simply evaluated through the indirect Lyapunov’s method; results are reported in Figure 1.

**Figure 1:**
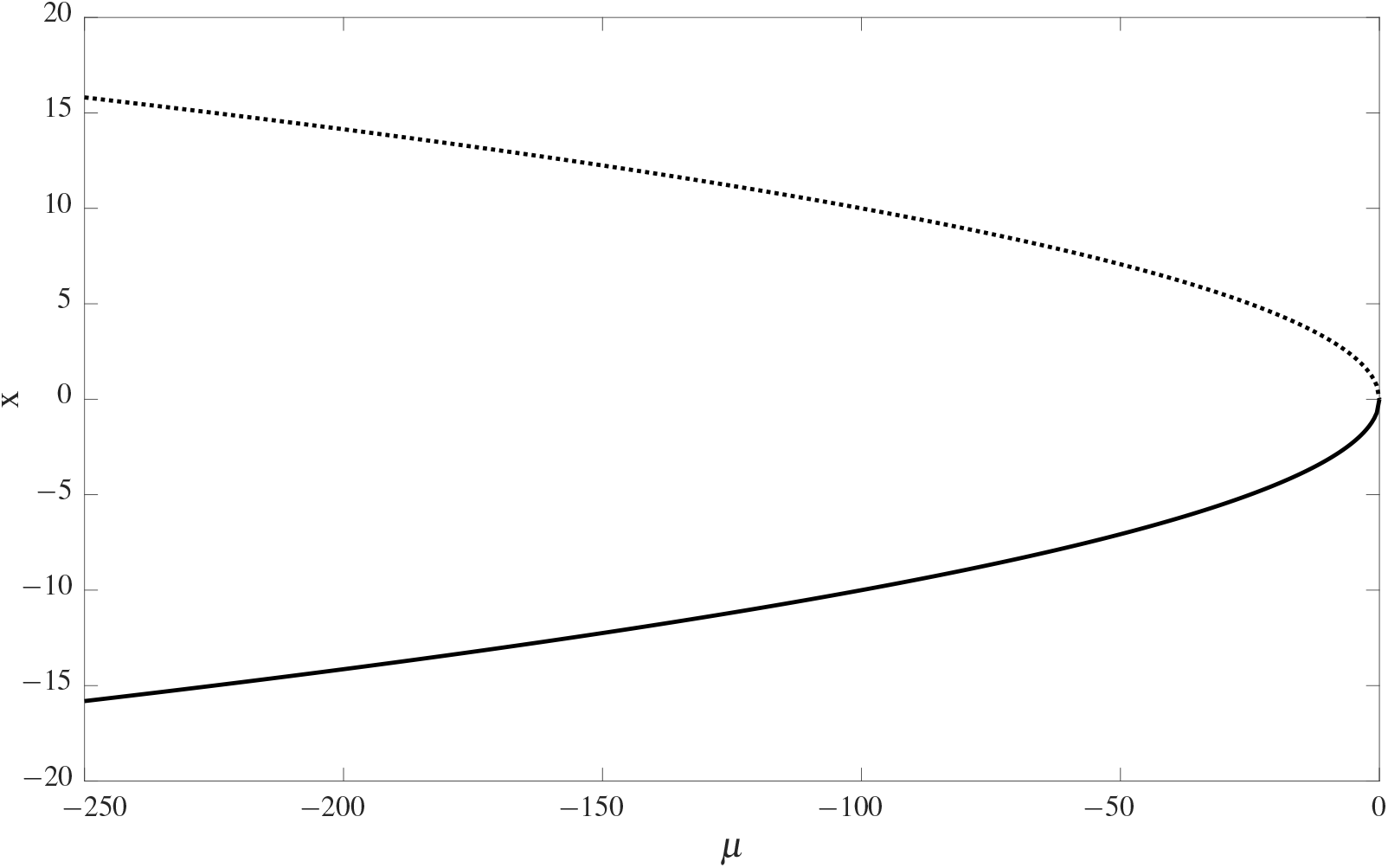
Fold bifurcation diagram. The position of the equilibria is reported against the value of the bifurcation parameter *μ*. Solid and dashed lines represent the stable and the unstable equilibria, respectively. The couple of equilibria, which coexists when the bifurcation parameter is negative, get closer when *μ* increases. When *μ* = 0 the two solutions collapse to the value *x* = 0, disappearing when *μ* > 0.

To trace out the equilibrium curve of (1), including both stable and unstable equilibria, CBC relies on the presence of a stabilising feedback controller. The equation of motion of the system including feedback control is given by

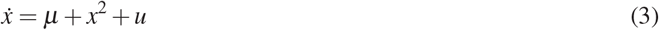

where *u* is the control signal given by a suitable, i.e. stabilizing, control law. CBC is not restricted to a particular form of control such that the control law can in principle take a general form:

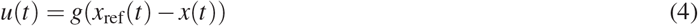

where *x*_ref_(*t*) is the control reference (or target). In this paper, simple linear proportional (P) and proportional-plus-derivative (PD) control laws will be considered such that *u* is given by

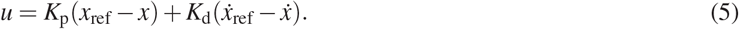

A PD controller is chosen due to its extreme simplicity and accessibility. Using (5) as control input, (3) becomes:

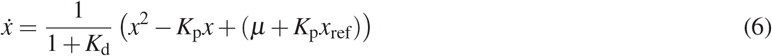

where *ẋ*_ref_ has been assumed to be equal to zero due to the focus on the equilibria of the system.

The equilibria of a controlled system are in general different from the one of the uncontrolled system. To recover the response of the underlying uncontrolled system of interest, CBC seeks a reference signal *x*_ref_ for which the controller is non-invasive [20], i.e. for which the control signal satisfies

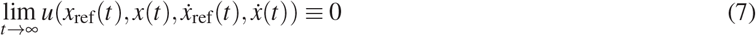

If (7) is satisfied, one can show that the equilibria of (3) are identical to those of (1). However, although the control signal is asymptotically zero, it changes the linearization of the dynamics such that unstable responses are now stable and hence observable. The search for the reference signal that leads to such a non-invasive controller can be performed using Newton-like algorithms.

Alternatively, when the control signal and bifurcation parameter affect the system in the same way (as in Eq. (3)), the static component of the control signal does not need to be removed and can be viewed as an additional contribution to the bifurcation parameter. Thus, we can write the equilibria of (6) and the control signal at steady state as:

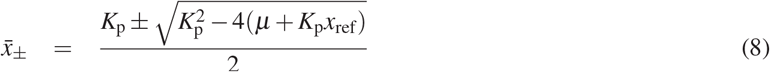

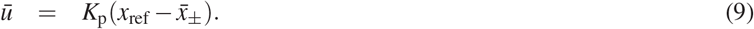

where *K_d_* ≠ −1 allowed a simplification of the equilibria definition in (6). Writing *x*_ref_ as function of the steady state control action *u*, we can substitute Eq. (9) in Eq. (8) as in the following passages:

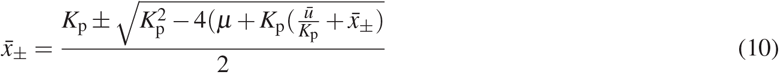

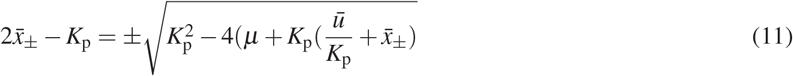

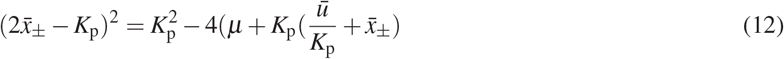

Solving the squared term and simplifying the equation we can thus obtain:

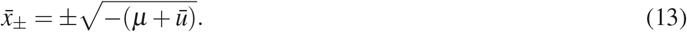

Eq. (13) defines the same equilibrium curve as Eq. (2) for a parameter *μ*\* = *μ* + *ū*. Therefore, the steady state contribution of the control signal does not have to be removed and it can simply be considered as a shift in parameter space. This result will also be exploited in the study of the pluripotency GRN in Section 2.3. Similar principles were also used to capture the periodic orbits and fold bifurcations of several mechanical systems [12–14, 21].

To capture unstable equilibria of system (1), the control gains *K*_p_ and *K*_d_ must be chosen such the controller has a stabilising effect. The stability of the steady-state response 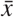 of (3) is govern by the sign of

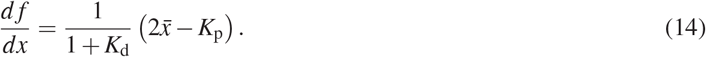

Using 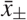 given by Eq. (8), we obtain

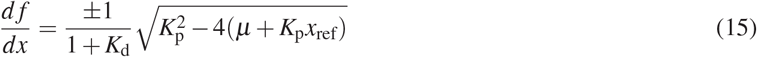

where the sign depends on the sign chosen for the equilibrium point 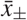. The equilibrium stability directly depends on the derivative gain value. In particular, if:

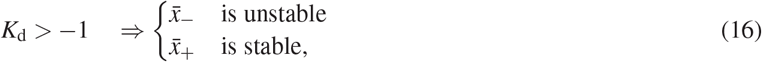

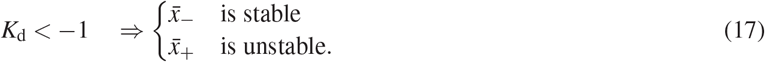

Furthermore, for Eq. (8) to have real solutions, it must be satisfied the following condition:

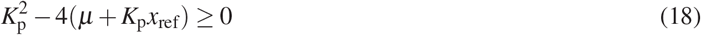

When (18) is not met, the system (3) does not have any equilibrium point.

Thus, combining Eq. (18) with Eqs. (16–17), four different scenarios arise. Figure 2 shows the position of the stable equilibria of the controlled system when the reference is varied. By fixing any one particular coupling of gains (*K*_p_, *K*_d_), it is not possible to evaluate all the equilibria of the uncontrolled system (1), but just a subset of them. It appears clear that it is possible to track all the equilibria of the system by opportunely changing the couple (*K*_p_, *K*_d_). The most convenient way to do so is to switch from *K*_p_ > 0, *K*_d_ > −1 to *K*_p_ < 0, *K*_d_ < −1 when the reference goes from negative to positive values, respectively.

**Figure 2:**
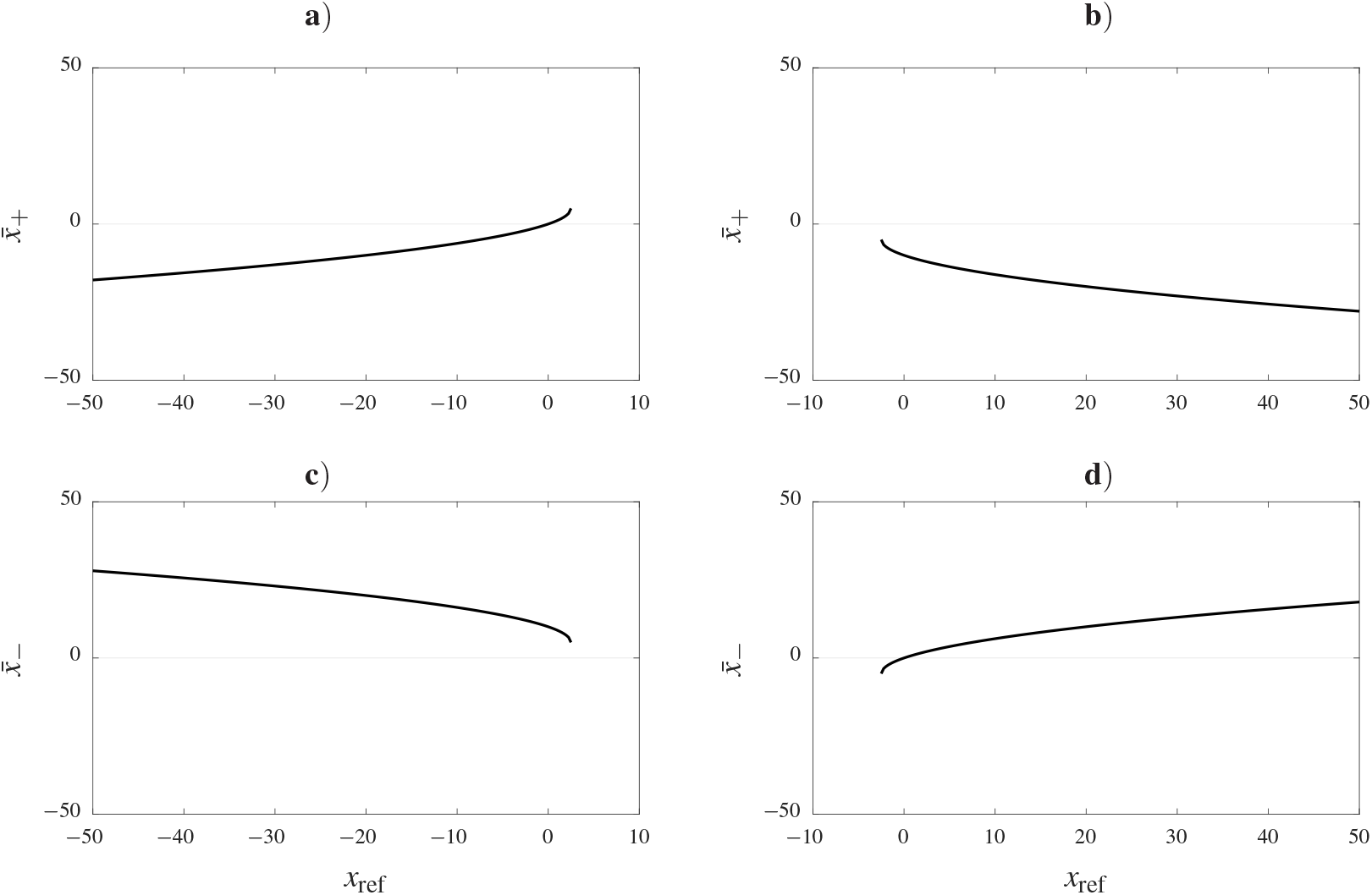
Controlled system stable equilibria in dependence of the reference signal (reported on *x* axis). Subfigures show respectively the cases: **a**) *K*_p_ > 0 *K*_d_ > −1, **b**) *K*_p_ < 0 *K*_d_ > −1, **c**) *K*_p_ > 0 *K*_d_ < −1, **d**) *K*_p_ < 0 *K*_d_ < −1.

### 2.2 Implementation of CBC

The practical implementation of the CBC method used in this paper can be summarized by the main following steps:

1. Under the assumption of a stabilizing control action, given a reference target *x*_ref_ the system is left to run for a fixed amount of time such that the transient dynamics dies out and the system reaches steady state. The achievement of a steady state solution can be verified by evaluating a condition on the values of the state and of the input. In particular, being:

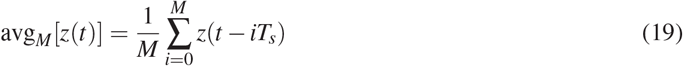

(where *z* stands for the quantity of interest, i.e. either *x* or *u*) if the difference between the averages of the control input and response signals obtained between two consecutive sample times are both below a user defined tolerance, the system is considered to have reached its steady-state. Otherwise, the system is left to run longer and the average values are check again after waiting.
2. At steady-state, the response and control signal are recorded, and the point 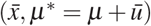 is an equilibrium point of the underlying uncontrolled system.
3. The algorithm moves on to capture a new point of the bifurcation diagram. To this end, the control reference target *x*_ref_ is stepped according to

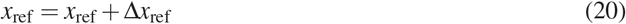

where Δ*x*_ref_ can either be a fixed value or adapted to satisfy a desired covering of the bifurcation diagram.
4. The algorithm is stopped when a desired number of points has been recorded or when the state or parameter have reached particular values.

The above algorithm has been applied to system (3) and the results are reported in Figure 3. The recorded data points always converge to a solution which lies on the analytical diagram of the fold bifurcation. Details of gains chosen are provided in the caption of Figure 3.

**Figure 3:**
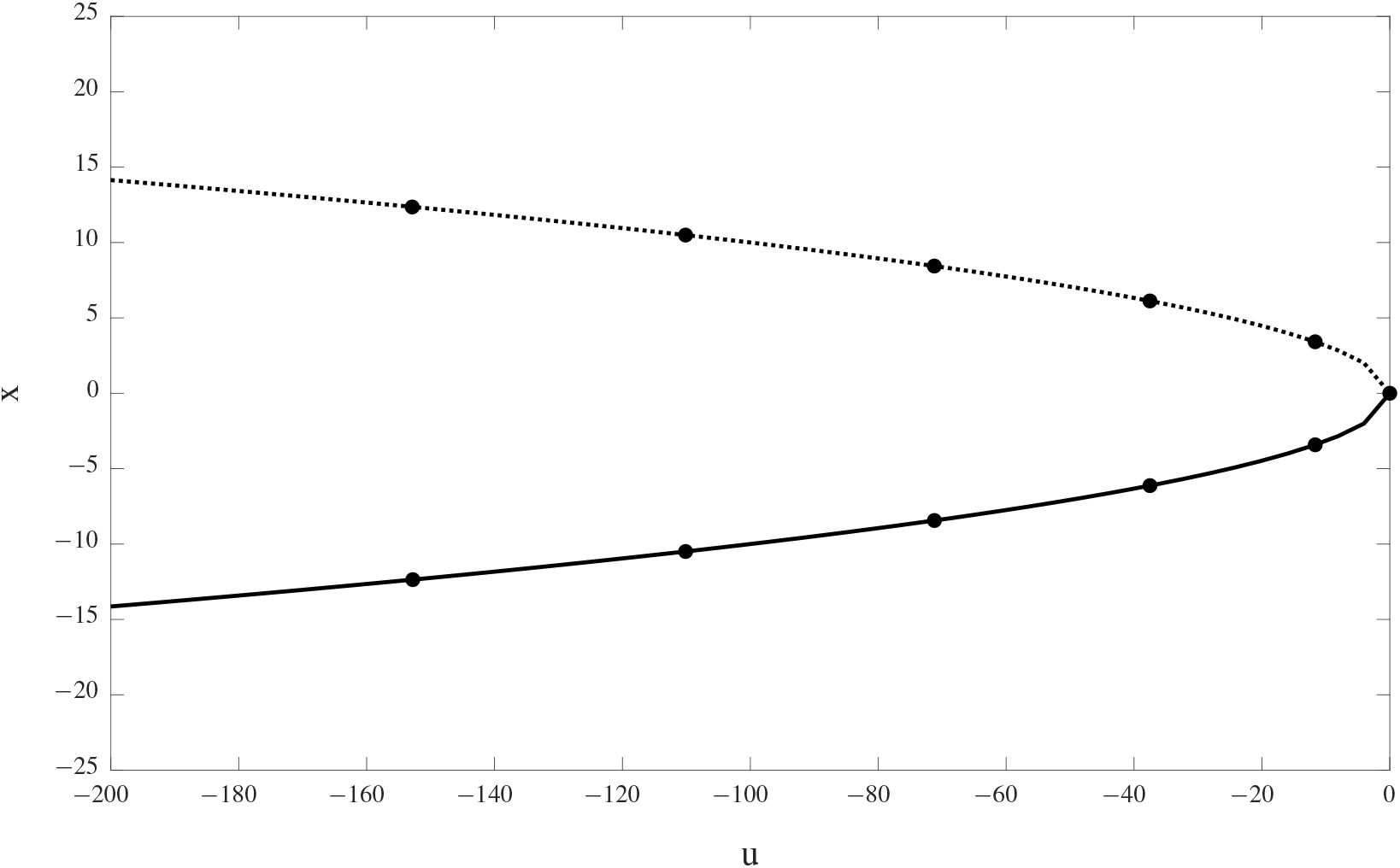
Results of the CBC applied to a fold bifurcation. Solid line represents the position, computed analytically, of the stable equilibrium for different values of the input, while dashed line represents the position of the unstable one. Bullet points are the equilibria computed using the CBC algorithm. Parameters of the algorithm: *x*_ref_ ∈ [−20,20], Δ*x*_ref_ = 4. The stabilizing control action is given by a PD controller whose gains are (*K*_p_, *K*_d_) = (20,2) when *x*_ref_ < 0 and (*K*_p_, *K*_d_) = (−20,−2) when *x*_ref_ ≥ 0.

### 2.3 Application of CBC to a pluripotency gene regulator

#### 2.3.1 Mathematical model of the pluripotency GRN

We now apply the method described in Sections 2.1 and 2.2, to track the equilibrium of a nonlinear biological system. A negative control action is physically meaningless, so for this implementation of the CBC we will forbid any negative value of *u*. The system of interest aims to capture the dynamics of a pluripotency GRN where Nanog, a master regulator of mouse embryonic stem cells pluripotency, has been reported experimentally to exhibit bistable steadystate accumulation in various culture media conditions [16–18]. In [19], Nanog heterogeneity has been associated to switch-like dynamics, with bistability caused by positive feedback loops in the underlying pluripotency GRN.

The dynamics of the pluripotency GRN are modelled using the following set of differential equations [19]:

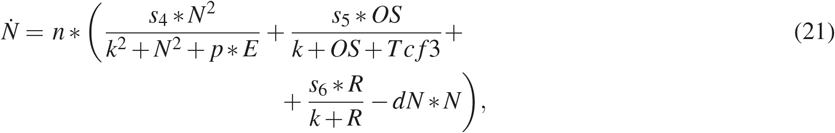

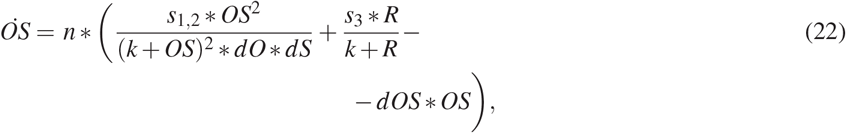

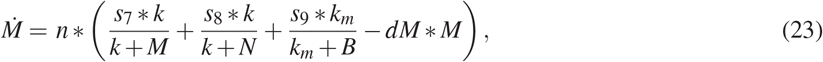

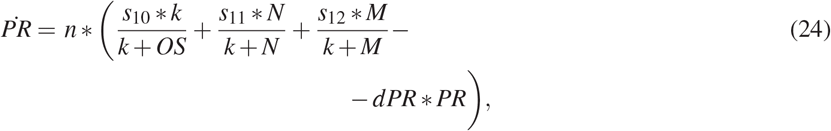

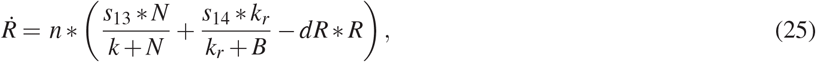

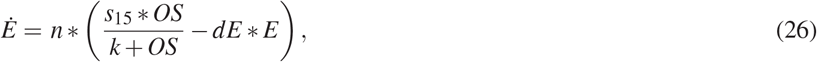

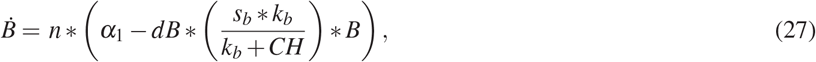

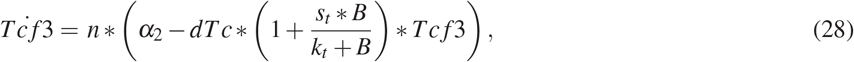

where the variables represent concentrations of the following genes or protein complexes: *N* is Nanog, *OS* is the Oct4-Sox2 complex, *M* is Mycn, *P* is Prdm14, *R* is Rest, *E* is FGF4/Erk, *B* is *β*-catenin. Regarding parameters, *n* is a scaling factor, *s_i_*(*i* = 1 – 15, *t, b*) are the maximal transcriptional rates, *k* and *k_i_*(*i* = *t, b, r, m*) are Michaelis-Menten constants, *d*[·] are degradation rates, *α_i_*(*i* = 1,2) represents basal activity; *CH* is the drug CHIR99021 (glycogen synthase kinase-3 inhibitor). *p* is a function of *Prdm*14 andthe MEK inhibitor PD0325901 *PD*, namely

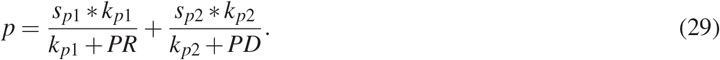

A comprehensive description of the model derivation and parameters, including their nominal values and units, is presented in [19].

We will be using *s*_4_, governing the strength of Nanog auto-activation, as the control input and bifurcation parameter for CBC, as in [19] we showed that variations in *s*_4_ can cause Nanog bistability. The latter result was also reproduced here: using the numerical continuation algorithm implemented in XPP/AUTO [22], we generated a full bifurcation curve, which shows the emergence of two saddle-node bifurcations separating two stable and one unstable Nanog solution (Figure 4a). Thus, in what follows, we aim at recreating the Nanog bifurcation diagram using CBC, having Nanog and (*s*_4_) as control output and control input/bifurcation parameter, respectively.

**Figure 4:**
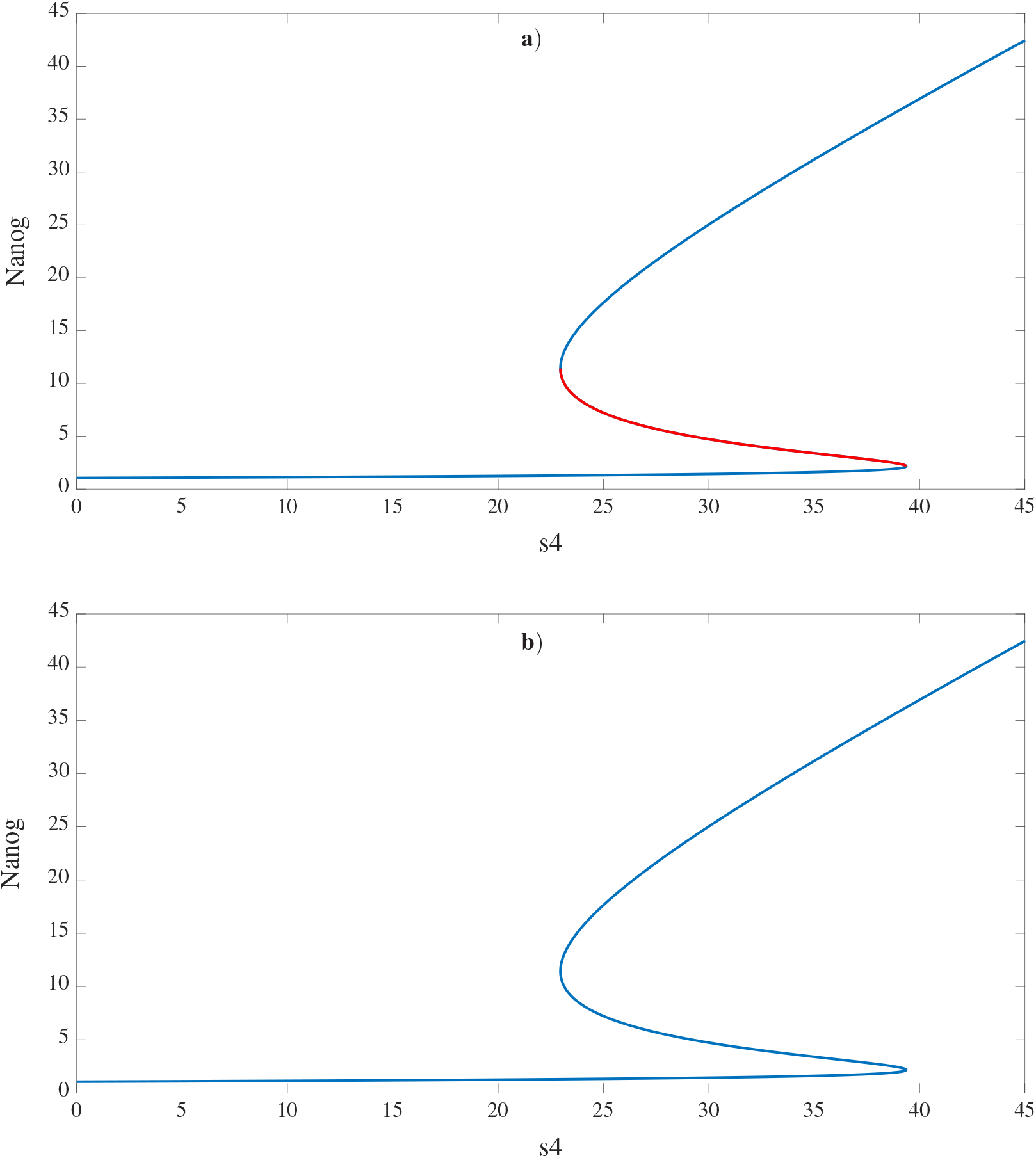
Numerical Continuation of Nanog system. Subfigure **a**): no control is applied. Red and blue lines denote unstable and stable Nanog solutions, respectively. Subfigure **b**): the Nanog system is subject to a proportional control with a proportional gain *K*_p_ = 5.95. Blue line denotes stable Nanog solutions. There are no unstable solutions in this case.

#### 2.3.2 Analytical study of Nanog equilibria stability for gain identification

We first performed a study to identify controller gains able to reproduce, with CBC, the numerical continuation results. To simplify the analysis and further show that CBC does not necessarily require sophisticated control strategies, we decided to use a proportional controller. In the model, we replaced the bifurcation and control parameter *s*_4_ (Equation (21)) with *s*_4_ + (*k_p_*(*x*_ref_ − *x*)), which implements our controller, where *x*_ref_ is the control reference and *k_p_* is the proportional gain, and we have renamed *N* to be *x* (i.e. the output).

We then evaluated the Jacobian matrix (*J*) of the system at Nanog steady state(*x*\*); *J* is an 8×8 matrix, and presented below are the terms in the Jacobian that are affected by the controller:

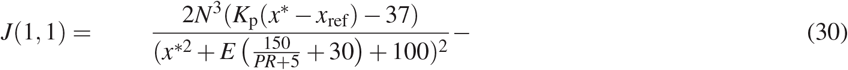

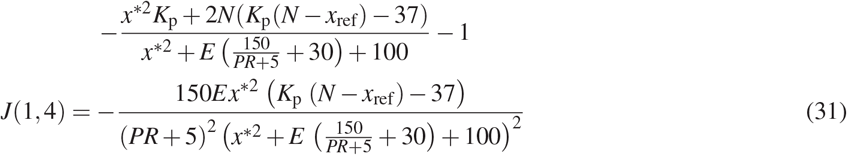

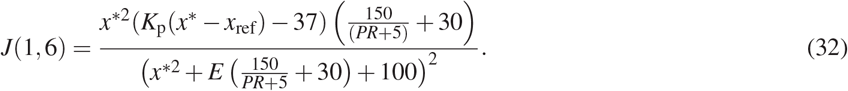

Using the Jacobian, we assessed the stability of Nanog steady state when varying the gain *K*_p_, with the aim of finding the smallest gain that is just over the critical point of being able to control the entire curve.

Figure 4 (where the stable and unstable solutions are denoted by blue and red lines, respectively) shows how the stability is affected: it is clear that, when increasing the strength of the controller, the region of instability shrinks, and finally disappears. The critical point for this is around *K*_p_ = 5.95, which is the minimum gain required for CBC to be able to stabilise the entire curve. Generally, a controller aims to reach steady state quickly. However, contributions to the control signal such as noise cannot be removed from the control signal, which means that using a controller with larger gains may lead to increased invasiveness, which is counter to the goal of CBC. Therefore, we will be using the critical value for the gain of the controller.

#### 2.3.3 CBC of Nanog dynamics

When performing CBC of the aforementioned system, we applied a constraint on the actuation time (i.e. the control action is applied every 60 minutes). This is due to the specifics of the control-actuation platform we foresee using when performing experiments in the future, where the control output is measured by means of fluorescent microscopy every 60 minutes to avoid photo-toxicity effects [23].

We used the numerical continuation curve to choose a maximum and minimum control reference point. The stepping method we used is constant but, as we said, it could also be made adaptive.

We used a rolling average of the last 5 samples taken, for both *u* and *x*, and a relative tolerance of 0.1% to satisfy our steady state check. When the rolling average of both the control input and system output have changed within the tolerance level, the reference point is stepped up.

Then, using the same method as in Section 2.1, we performed CBC on the Nanog system (Figures 5b and 6). The experiment was a simulation of slightly over 41 days for 41 points and was successful in accurately capturing the dynamics of the system, as the points follow the numerical continuation curve of the system model. Of note, the corresponding brute force continuation of Nanog steady state by varying *s*_4_ (Figure 5a), using a MATLAB solver for a range of *s*_4_ values, shows the limitations of an open loop approach, as it cannot capture the unstable steady states of the system. Indeed, any simulation that is run around the *s*_4_ values corresponding to the unstable steady states jumps, as expected, to one of the two stable equilibria. Instead, CBC can track the full curve, as shown in Figure 5b, and precisely capture all the steady states of the system through the use of a suitable non-invasive controller.

**Figure 5:**
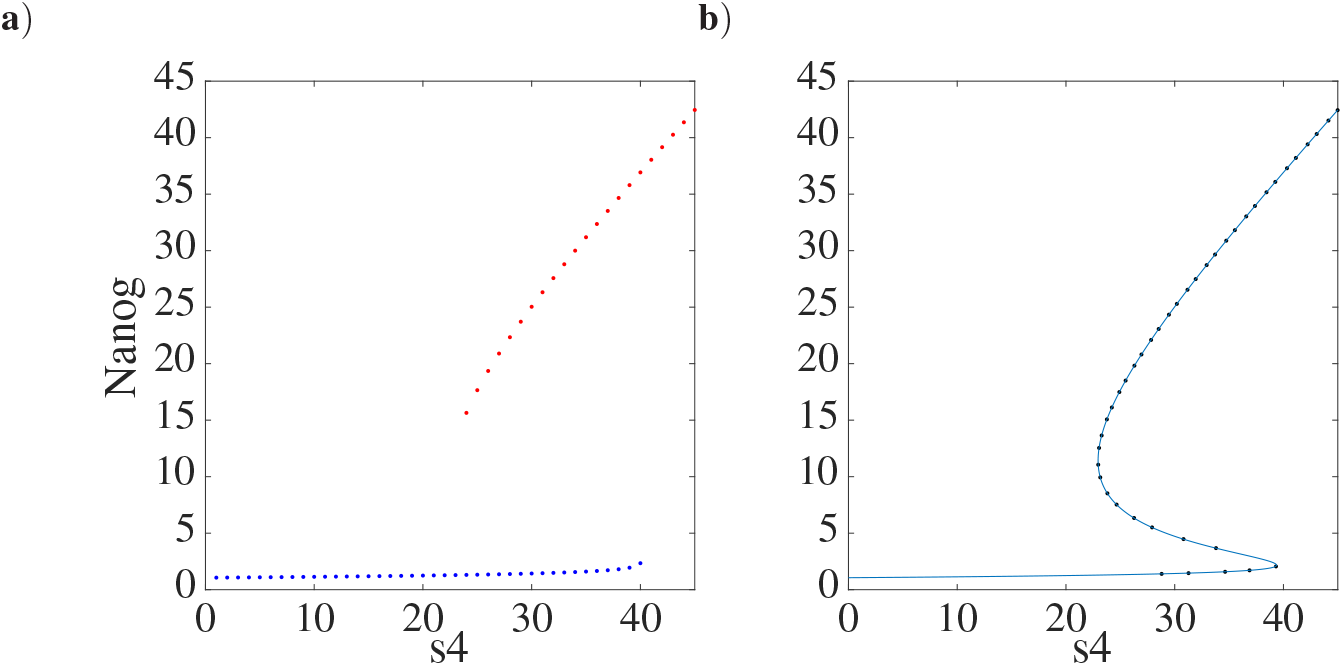
**a**) Brute force continuation of Nanog. In red, we ran a brute force simulation starting from high values of *s*_4_ and decreasing them. In blue, we started from low values of *s*_4_ and increased them. **b**) CBC of Nanog dynamics using a fixed stepping reference. The black dots indicate the CBC result and the blue line represents the previously done numerical continuation of the system.

**Figure 6:**
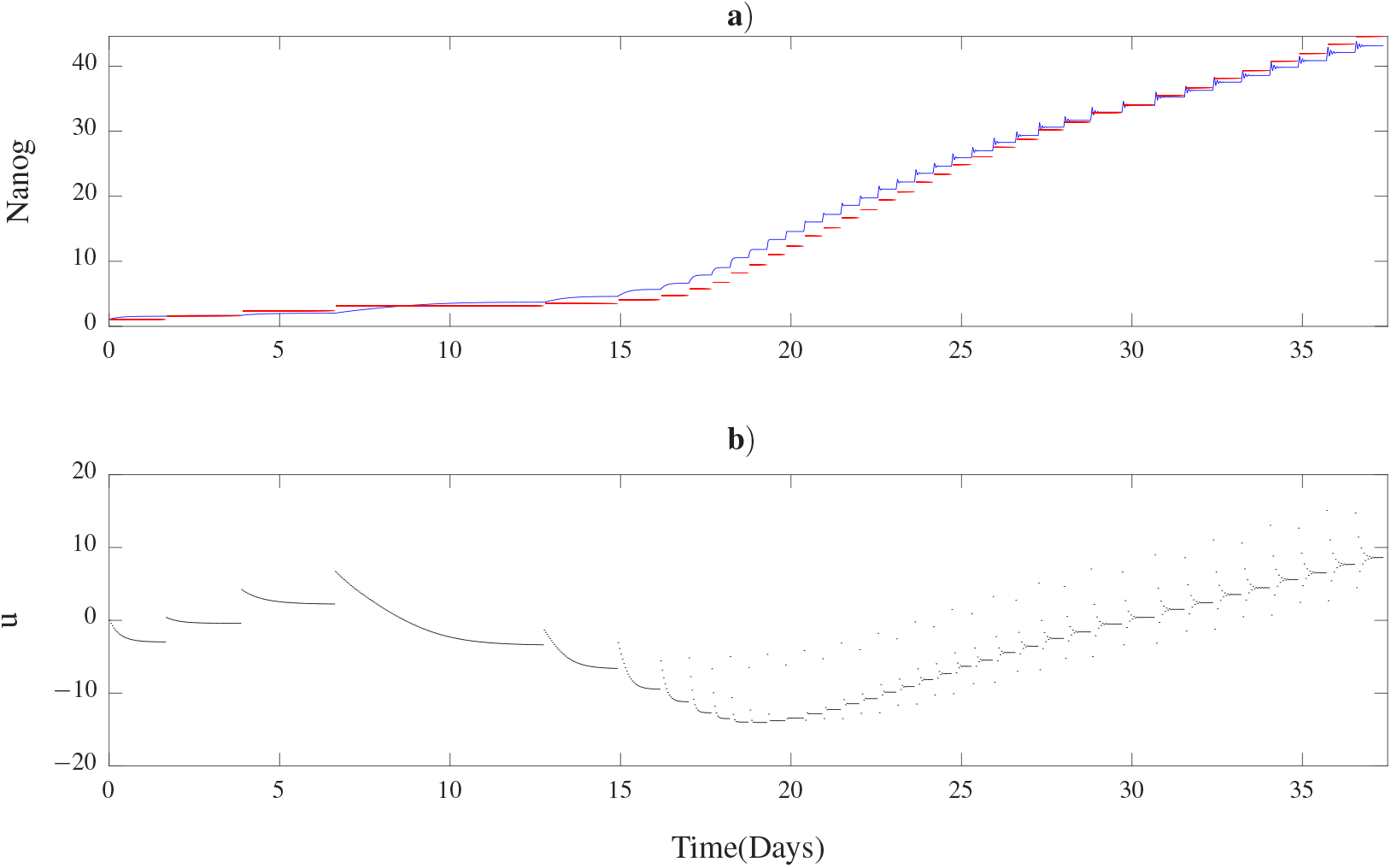
Plots of the output and control input of the system over the time of the experiment. **a**) Graph of Nanog output against time; red points represent the given reference, blue ones represent the output of the system. **b**) Graph of control input against time; black points represent the control input.

In Figure 6 we can see that there is a wide variety in the amount of time that it takes for a data point to satisfy the conditions to move to the next reference point. It is worth taking into consideration that the data points that take the longest time to converge are those at the lower end of the CBC curve (Figure 6); this might be due to the fact that, when the curve has a slow ascending gradient, it takes longer for the controller to enable the output to reach a unique value. This issue could be solved by using a stronger controller. However, for points that are higher on the curve, we can see that the controller we used might be too strong, as the input fluctuates between positive and negative values and causes some small oscillations in the output before the transient dynamics of both controller and output have been nullified to a suitable degree for the algorithm to continue (Figure 6). We foresee that an adaptive controller, which can reduce the oscillations and increase the rate of convergence at the upper and lower end of the curve, respectively, could address these issues.

Of note, we do not require the output of the system to precisely reach the reference point, as shown in Figure 7 (i.e., a magnification for a single point of the CBC simulated experiment in Figure 6). Indeed, we only require the controller to make the output reach a unique value that differs from the previous value obtained, as the control target is just a proxy for the states. If the output were to match the reference point exactly, the algorithm would set the control to be zero; this, in turn, would prevent CBC from obtaining data points at different parameter values (as the control signal represents a shift in parameter space).

**Figure 7:**
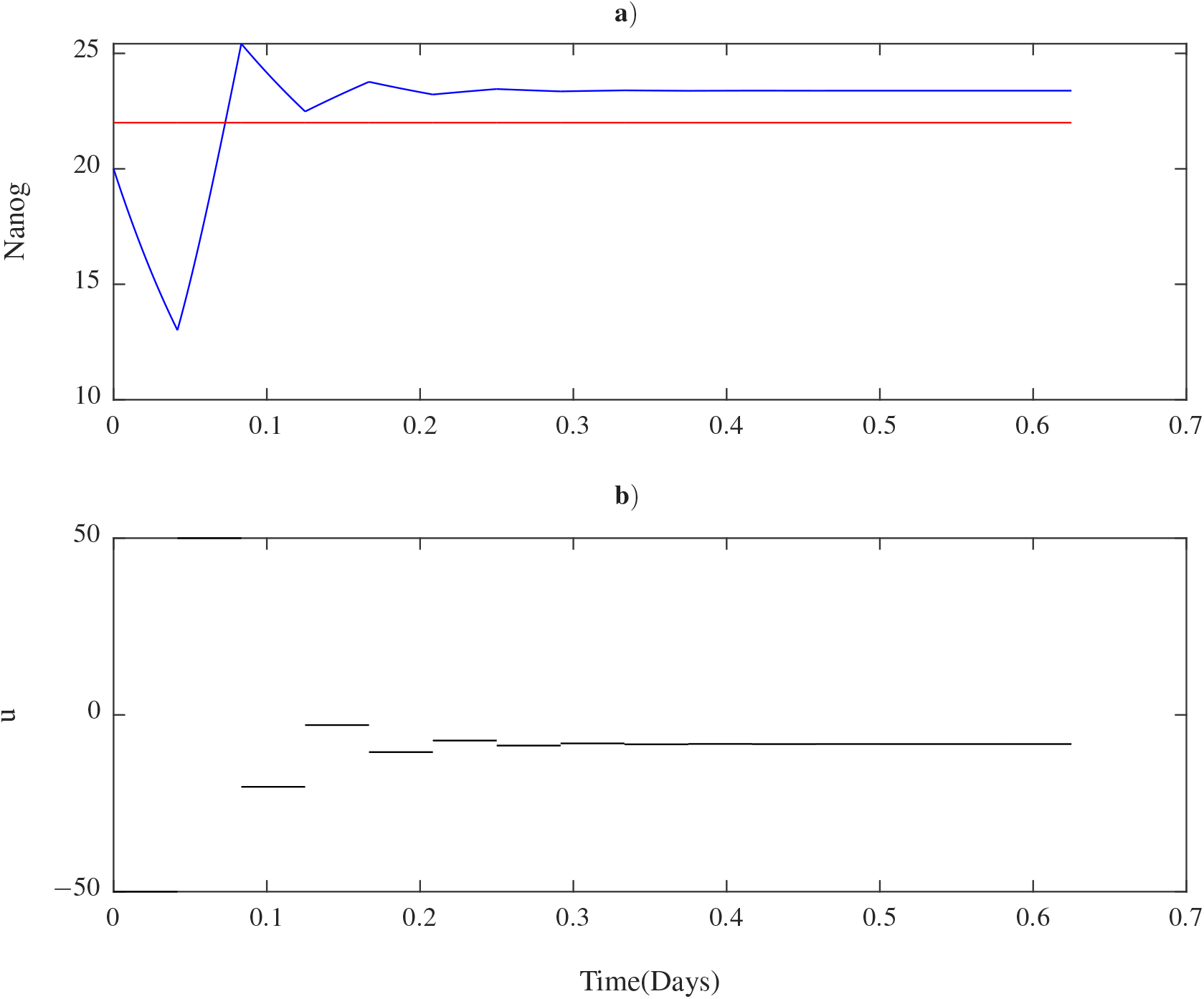
The control action implemented on a single point.**a**) Graph of Nanog output against time: red points represent the given reference, blue ones represent the output of the system. **b**) Graph of control input against time: black points represent the control input used.

## 3 CONCLUSIONS

In this work we provided the first proof of the applicability of CBC for tracking solutions of biochemical systems. The next steps in this research will involve further *in silico* studies including noise in the system dynamics to better represent the stochastic nature of biochemical interactions, and implementing CBC experiments to directly measure and control Nanog in living mouse embryonic stem cells by means of fluorescent reporters, and microfluidics/microscopy platforms for external feedback control of Nanog expression [23,24]. We believe CBC could be extended to Synthetic Biology applications to enable model-free prototyping of engineered GRN with desired nonlinear dynamics.

## ACKNOWLEDGMENT

This work was supported by the COSY-BIO project, which has received funding from the European Unions Horizon 2020 under Grant Agreement No. 766840 to M.d.B. and L.M. (and related Scholarship to A.G.), by an Engineering and Physical Sciences Research Council PhD Scholarship to B.G., by the Royal Academy of Engineering Research Fellowship (RF1516/15/11) to L.R., by the Medical Research Council grant MR/N021444/1 to L.M., by the Engineering and Physical Sciences Research Council grant EP/R041695/1 to L.M., and by BrisSynBio (a BBSRC/EPSRC Synthetic Biology Research Centre) BB/L01386X/1 to L.M. and MdB.

